# Murine trypanosomiasis recapitulates transcriptomic features of acute kidney injury

**DOI:** 10.1101/2024.05.08.593024

**Authors:** John Ogunsola, Anneli Cooper, Juan F. Quintana, Annette MacLeod

## Abstract

The African trypanosome, *Trypanosoma brucei,* disseminates systemically in tissues of the infected host resulting in complex immunopathology. The kidneys which are important in the response to the anaemia characteristic of African trypanosomiasis, are prone to acute kidney injury (AKI) from multiple noxious stimuli. Little is known about the transcriptional responses of the kidney to trypanosome infection. To assess the tissue-specific response to infection with *Trypanosoma brucei*, we profiled the clinicopathologic and transcriptional responses of the kidney in BALB/C (susceptible) and C57BL/6 (tolerant) murine models, at early (7 dpi) and late (21 dpi) time points of infection. Trypanosomes in the renal interstitium, tubular necrosis and inflammation characterised early infection in both mouse strains. By late infection, we observed extensive tubular necrosis in the susceptible BALB/C but reparative tubular regeneration in the tolerant C57BL/6 mice. *T.b. brucei* infection resulted in significant increases in serum creatinine in both strains. Consistent with the clinicopathologic findings, RNA-seq detected both mouse strain- and time-dependent transcriptional responses in the kidney. These included perturbations in genes associated with solute/ion transport, upregulation of markers of tubular injury, hypoxia, glycolysis, and a profound inflammatory and immune response, mirroring the responses observed in other models of AKI. Differential tissue pathology at late time point is preceded by expansion of CD8^+^ T cells, profound expression of transcription factors and upregulation of anti-inflammatory pathways in C57BL/6 mice. Our findings demonstrate that experimental *T. brucei* infection-induced kidney injury (TIKI) is a model of AKI and may have clinical implications for Human African Trypanosomiasis cases, who currently are not routinely screened for markers of kidney function.

## Introduction

The African trypanosome, *Trypanosoma brucei,* is of immense economic and medical importance in sub-Saharan Africa where it causes potentially fatal disease in humans and domestic animals, through Human African Trypanosomiasis (HAT) and Animal African Trypanosomiasis (AAT), respectively. These flagellated protozoan parasites are transmitted by a tsetse vector (*Glossina spp*) (Vickerman 1985) and cause an initial haemo-lymphatic infection followed by a later meningo-encephalitic stage when trypanosomes colonise the central nervous system (Kennedy 2013). Historically, *T. brucei* has been regarded as a blood- and lymph-dwelling parasite, but recent reviews have challenged this view with colonisation of several tissues described (Alfituri *et al*. 2020; Crilly and Mugnier 2021). In humans, domestic animals and experimental models, extravascular compartments such as skin (Capewell *et al*. 2016; Camara et al. 2020; Quintana *et al*. 2023), circumventricular organs (Quintana *et al*. 2022), adipose (Trindade *et al*. 2016; Machado *et al*. 2021; Sinton *et al*. 2023) and reproductive organs (Carvalho *et al*. 2018) have been characterised.

*T. brucei* survives and multiplies in tissue compartments of their mammalian host despite being confronted by the innate and adaptive arms of the immune system. In visceral organs and the associated microenvironmental niches, host-derived factors are thought to impact on parasite survival, growth dynamics and transmission (De Niz *et al*. 2021). A combination of parasites’ tissue colonisation and the associated host’s inflammatory and immune responses also result in well described organ-specific pathologies such as lymphadenopathy, hepatosplenomegaly, orchitis and pancarditis (Stijlemans *et al*. 2016). These pathologies mediate clinical signs of the disease.

One of the main clinical features of trypanosome infection is anaemia (Brun and Blum 2012). In response to anaemia and tissue hypoxia that ensues, the kidneys secrete the hormone erythropoietin which drives production of red blood cells (Moore and Bellomo 2011). The functional unit of the mammalian kidney is the nephron which comprises of a glomerulus and an extensive, multi-segment tubule (Chi *et al*. 2006). Due to the anatomic peculiarities of its blood supply, intense oxygen requirements and extreme reliance on aerobic respiration, the kidneys are sensitive to hypoxia and prone to acute kidney injury (AKI) (Evans *et al*. 2020). AKI is characterized clinically by abrupt reduction in urine output and elevation of serum creatinine and results in significant morbidity if left uncontrolled (Liu *et al*. 2017). Hypoxia is one of the notable causes of AKI. Despite multiple causes of AKI, a common pathologic feature is tubular injury (Gaut and Liapis 2021), thus suggesting a unified tissue response at the transcriptomic level. At least four mouse models to induce AKI (including ischemia-reperfusion, sepsis, malignant hypertension, rhabdomyolysis, and cisplatin toxicity) have been developed and they share similar molecular characteristics and outcomes (Hultström *et al*. 2018). However, our understanding of the pathology and molecular response of the kidneys during *T. brucei* infection is limited.

Severity of organ pathologies, following *T. brucei* infection in susceptible hosts, is in part due to host genetics (Naessens 2006; Noyes *et al*. 2009). In mice, ‘trypanotolerance’ characterized by mild disease is linked to a control of the parasitaemia (Naessens 2006). In murine models, different strains have been observed to demonstrate differing degrees of susceptibilities to trypanosome infection (Magez *et al*. 2004). Here, using two murine models with differential susceptibilities to trypanosome infection, RNA sequencing and informatic approaches, we set out to characterise the clinicopathological, immunological and transcriptional responses of the murine kidney to acute and chronic *T. brucei* infection.

Our studies demonstrate that *T. brucei* infection induces histological and transcriptional responses comparable to those observed in other models of AKI. Furthermore, our results demonstrate that in the trypanotolerant C57BL/6 mouse model, the trypanosome infection-induced kidney injury (TIKI) is resolved, likely mediated by the upregulation of tissue repair gene pathways, a controlled upregulation of CD8+ T cell-associated transcripts and a superior response to hypoxia, whereas the trypanosusceptible BALB/c mice maintain kidney pathology driven by markers of T cell exhaustion such as PD-1/PD1L. Together, our findings demonstrate that experimental *T. brucei* infection results in infection-induced AKI, and presents an attractive model to explore renal responses to infection. These observations have clinical implications for HAT patients who currently are not routinely screened for markers of kidney function.

## Materials and methods

### Ethics statement

All animal experiments were approved by the University of Glasgow Ethical Review Committee and performed in accordance with the Home Office guidelines UK Animals (Scientific Procedures) Act, 1986 and EU directive 2010/63/EU. All experiments were conducted under SAPO regulations and UK Home Office project licence number PC8C3B25C.

### Trypanosoma brucei infection

Eight-week-old, female C57BL/6 mice (*n* = 7) and BALB/c mice (*n* = 6) (JAX, stock 000664) were inoculated by intraperitoneal injection with 3,000 parasites of strain *T. b. brucei* Antat 1.1E (Le Ray *et al*. 1977). Parasitaemia was monitored by regular sampling from tail venepuncture and examined using light microscopy and the rapid “matching” method (Herbert and Lumsden 1976). Uninfected mice of the same strain, sex and age, served as uninfected controls (*n* = 3). Mice were fed *ad libitum* and kept on a 12-hour light-day cycle. Samples were collected to capture the histologic and transcriptional changes in the murine kidney during the first peak of infection at 7 days post infection (dpi) and during late stage of infection at 21 dpi. Both kidneys were removed from each mouse; the left kidney preserved in RNAlater (ThermoFisher) for RNA extraction and the right kidney preserved in neutral buffered formalin for histopathology analysis.

### Histopathology scoring of the murine kidneys

Kidney slices were fixed in formalin for 24 hours at 4°C before proceeding with paraffin-embedding. 3 µm-thick histologic sections were stained with routine haematoxylin and eosin (H&E) or periodic acid Schiff (PAS) and mounted. Microscopic examination of the sections was performed with the pathologist blinded to the infection time point and mouse strain. A semi-quantitative grading scale for the severity of lesions (with 0, 1, 2 and 3 corresponding to absent, mild, moderate, and marked respectively), was designed to evaluate specific pathologies of the glomeruli, tubules and renal interstitium. For each section, 20 glomeruli were counted and the number of glomeruli with >3 mesangial cells per mesangial area was determined: <1, 1-5, 6-10, and >10 glomeruli corresponded to absent, mild, moderate, and marked respectively. The tubular compartment in 20 random high-power fields of the renal cortex was examined for evidence of tubular degeneration (vacuolation, loss of brush border) and necrosis (pyknosis, karyorrhexis). Severity of tubular lesions were graded based on the number of high-power fields where lesions were present: 0-1, absent; 2-4 mild; 5-7, moderate and >7 marked. The number of foci of inflammatory aggregates in the renal interstitium were quantified and graded: 0, absent, 1-2 foci as mild, 3-5 foci as moderate and >5 foci as marked. A composite score (sum of the score for each compartment) was generated for each mouse to compare the severity of renal lesions.

### Immunohistochemistry

Immunohistochemistry was performed to detect and demonstrate African trypanosomes in the murine kidney. Rabbit anti-trypanosome BIP diluted at 1:10,000 was used, kindly donated by JD Bangs University at Buffalo,. Sections were counter- stained with haematoxylin.

### *Trypanosoma brucei* quantitation in mice kidneys

To detect trypanosome DNA in the kidneys, the number of copies of the *Pfr2* gene present within 20 ng of DNA prepared from approximately 30 mg of kidney homogenate was determined by quantitative TaqMan PCR. A modified protocol as described by Laperchia *et al*. (2016) was employed. Briefly, TaqMan PCR, using primers and probe specifically designed to detect the trypanosome *Pfr2* gene, was performed in a 25 μL reaction mix comprising 1× TaqMan Brilliant II master mix (Agilent, UK), 0.05 pmol/μL forward primer (CCAACCGTGTGTTTCCTCCT), 0.05 pmol/μL reverse primer (GAAAAGGTGTCAAACTACTGCCG), 0.1pmol/μL probe (FAM-CTTGTCTTCTCCTTTTTTGTCTCTTTCCCCCT-TAMRA) (Eurofins, Germany) and 20 ng template DNA. A standard curve was constructed using a serial dilution (range; 1 × 10^6^ to 1 × 10^2^ copies) of pCR®2.1 vector containing the cloned *Pfr2* target sequence (Eurofins). The amplification was performed on a MxPro 3005 (Agilent) with a thermal profile of 95°C for 10 minutes followed by 45 cycles of 95°C for 15 seconds, 60°C for 1 minute and 72°C for 1 second. The MxPro qPCR software (Agilent) was used to generate a standard curve and extrapolate the relative preponderance of trypanosomes using the number of Pfr2 copies as a proxy.

### RNA extraction and transcriptomics analysis

Half of the kidney (approximately 30mg) was transferred into an RNAse-free tube containing a lysing Matrix M bead (MPbiomedical). Tissue was lysed in a mechanical tissue lyser (Qiagen LT) at 50 revolutions per second for 2 minutes. From this lysate, RNA was extracted using the RNeasy Mini Kit (Qiagen) according to the manufacturer’s instructions. RIN scores assessed using a Bioanalyzer (Agilent; 2100 Expert) were >7. All RNA samples were submitted to the Beijing Genomics Institute (BGI, Hong Kong) for library preparation, sequencing, and quality control. Briefly, prior to sequencing, mRNAs were enriched using oligo-dT beads. This was followed by fragmentation, first-strand and second-strand cDNA synthesis. The synthesized cDNA was subjected to end-repair and then was 3’ adenylated. Adaptors were ligated to the ends of these 3’ adenylated cDNA fragments. DNA nanoball synthesis followed rounds of PCR amplification of the cDNA. Sequencing was performed on the DNBSEQ Technology platform to generate 150bp paired-end reads. The total number of raw reads is presented in **Supplementary S1**. Filtered, high-quality reads were aligned to the *Mus musculus* genome (GCF_000001635.26_GRCm38.p6) (Zerbino *et al*. 2017) using HISAT2 (Kim *et al*. 2015) with default parameters. To quantify expression, Bowtie2 (Langmead and Salzberg 2012) was used to map the clean reads to the reference gene sequence (transcriptome), and then RSEM (Li and Dewey 2011) to calculate the gene expression level of each sample. *In silico* analysis for deconvolution of inflammatory cellular composition in each sample was performed using seq-ImmuCC (Chen *et al*. 2017). Differential gene expression analysis was performed using DESeq2 (v1.18.1) (Love *et al*. 2014). Genes having an adjusted *p*-value < 0.05 and a log2fold-change (LFC) < -0.5 or > 0.5, across comparisons were considered differentially expressed. Comparisons were performed for each mouse strain: naïve versus early infection, naïve versus late infection, and early infection versus late infection. Gene lists for each comparison for each mouse strain were generated. Raw and processed RNA-sequencing data files have been deposited in the GEO database under the accession number GSE224927. Venn diagrams, heatmaps and PCA plot were created using package *ggplot2* (v3.0.0) (Wickham 2016) in the RStudio (v2021.09.0) interface of the R (v3.6.3) statistical software.

### Gene Ontology Analyses

To understand renal-specific transcriptomic responses, we adopted a bioinformatic approach where we compared our dataset to other published reports/datasets. Specifically, we compared the lists of dysregulated genes for both naïve-versus-early and naïve-versus-late comparisons in both mouse strains to previously published mouse atlases of kidney compartment-specific genes (Lee *et al*. 2015; Park *et al*. 2018). We also compared our dataset to a previously reported list of genes common to 4 models of AKI in humans (Hultström *et al*. 2018). We performed Gene Ontology Biological Process annotation (Ashburner *et al*. 2000) on downregulated gene sets for each comparison using the web-based *g:Profiler* (version e106_eg53_p16_65fcd97) (Raudvere *et al*. 2019) with the following settings: *i*) an ordered query of dysregulated genes, *ii*) Benjamini correction method with significance threshold of 0.05, *iii*) the statistical domain comprised only of annotated genes and term sizes ranged from a minimum of 5 to maximum of 500. We then focused on a subset of genes with log2fold- change (LFC) > 1, from our upregulated gene lists for each comparison. Each upregulated (LFC > 1) subset gene list was put through *g:Profiler* using settings as outlined above to determine either Kyoto Encyclopaedia of Genes and Genomes (KEGG) (Kanehisa et al., 2002) pathways. Finally, we then focused on common differentially expressed genes that were dysregulated throughout infection (that is, genes in the intersection of naïve-versus-early (NvE), naïve-versus-late (NvL) and early-versus-late time point (EvL) comparisons) for each mouse strain. Within the intersection gene list for each strain, we inputted genes that were upregulated for all comparisons into *g:Profiler* and performed KEGG pathway enrichment. Similarities and differences among the enriched KEGG pathways in both mouse strains were noted.

### Serum creatinine measurements

#### Mouse

Sera was obtained from each mouse at experimental endpoints. Serum creatinine was measured using the *Abcam* serum creatinine ELISA kit (ab65340) in accordance with manufacturer’s instructions.

#### Human subjects

Sera was obtained from the TrypanoGEN biobank (Ilboudo *et al*. 2017) of individuals with gambiense HAT in the DRC. Plasma creatinine was measured in freshly thawed plasma samples using the CREP2 kit (Catalog Number 03263991190) on a Cobas 311 analyser (Roche, USA). Assays were performed by a commercial laboratory in the UK. Samples were de-identified except for their HAT status (controls or cases) and necessary approvals from ethical boards received.

### Statistical Analysis

Parasite load in the blood (parasitaemia) and kidney (*Pfr2* gene copies) are presented as mean ± SEM. An independent *t* test or the non-parametric Mann-Whitney U-test (for ordinal values) was performed to compare the histopathologic scores between any two groups in a strain- or time-dependent manner. For more than two groups, a Kruskal-Wallis test was used. A *p*-value <0.05 was considered statistically significant.

## Results

### *Trypanosoma brucei* colonises the kidney early in infection resulting in clinicopathological features of AKI

Here, we set out to characterise the responses of the kidney to *T. brucei* infection using two murine models of infection, BALB/c and C57BL/6 mice, considered to be susceptible and tolerant to the infection by African trypanosomes, respectively (**Figure 1A**). In both strains, mice developed detectable parasitaemia by 4 days post infection (dpi) which reached first peak at 6 dpi and was not significantly different between BALB/c (2 × 10^8^ parasites/ml) and C57BL/6 (1 × 10^8^ parasites/ml). Following this initial peak parasitaemia, the strains diverged in their abilities to control parasitaemia. Whereas the trypanotolerant C57BL/6 mice successfully controlled parasitaemia to less than 10^6^ parasites/ml following the first (6 dpi) and second (14 dpi) peaks of infection, the parasitaemia remained high in the trypanosusceptible BALB/c mice for the duration of the infection (**Figure 1B**). Our findings are consistent with other studies (Magez *et al*. 2004; Magez and Caljon 2011) that describe the parasitaemia in these strains of mice with differential susceptibility to *T.b. brucei* infection with C57BL/6 being relatively trypanotolerant and BALB/c mice as trypanosusceptible.

**Figure 1.**
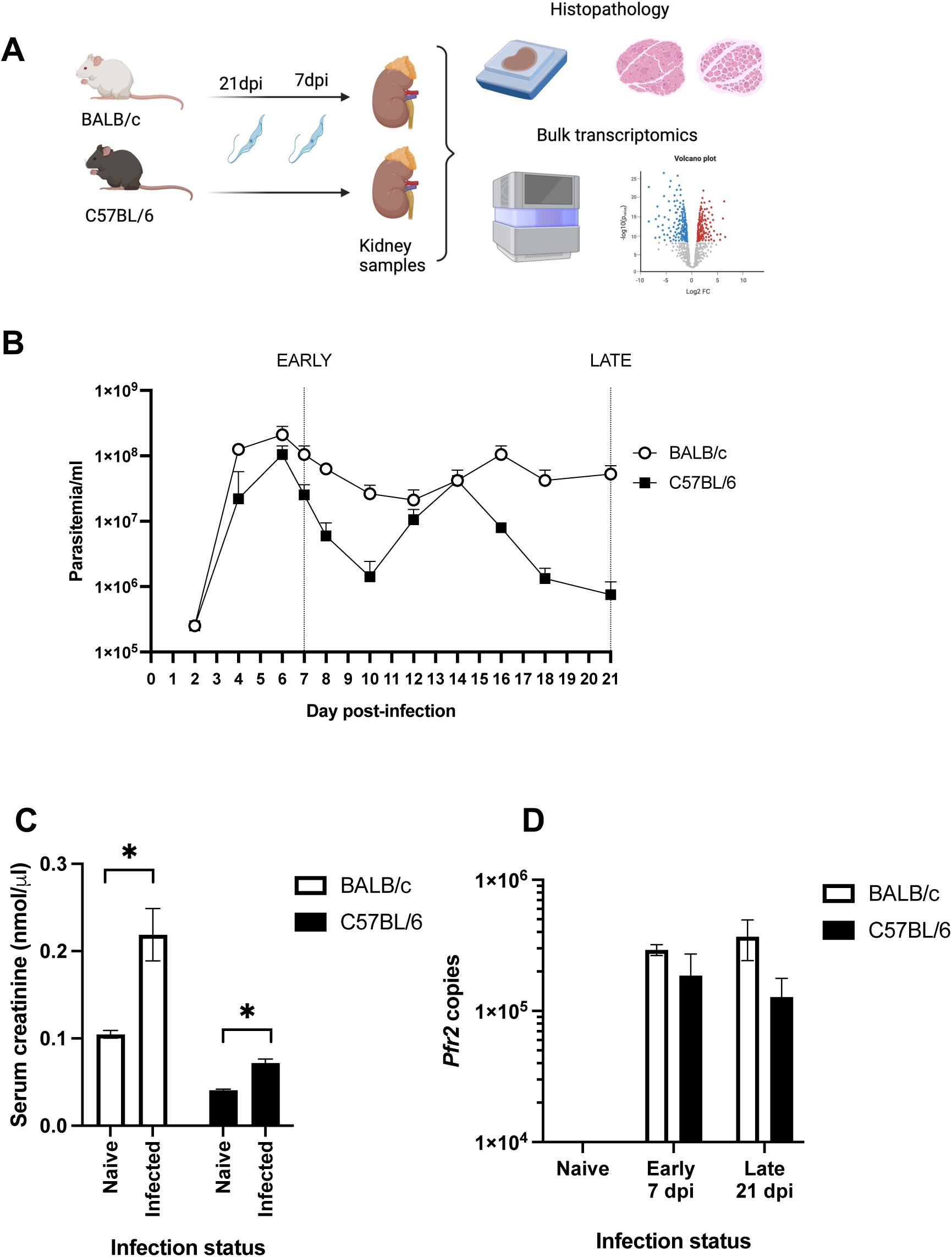
Susceptible BALB/c and tolerant C57BL/6 mouse models of *Trypanosoma brucei* infection. **A)** Experimental design and workflow. Susceptible BALB/c and tolerant C57BL/6 mice were infected with *Trypanosoma brucei* and monitored till day 21 post infection. Kidneys were extracted at early (7 dpi) and late (21dpi) time points for histopathology and bulk transcriptomics. Differentially expressed genes within several comparisons were either compared with pre-published gene lists or analysed for functional pathway enrichment using the KEGG database. **B**) Parasitaemia. *Trypanosoma brucei*-infected tolerant (C57BL/6) and susceptible (BALB/c) mice show comparable first peak of parasitaemia but control parasitaemia differently during infection. **C)** Serum creatinine levels of susceptible BALB/c and tolerant C57BL/6 naïve mice compared to *T. brucei* infected mice (both early and late timepoints). * *p* < 0.05. **D)** Parasite burden in the kidney. Estimation of *Trypanosoma brucei* DNA in the kidneys of tolerant (C57BL/6) and susceptible (BALB/c) mice using the parasite Pfr2 gene as proxy.

To assess kidney function we measured serum creatinine in both strains to determine the effect of infection. The early and late timepoints were combined as a single *T.b. brucei* infection group to increase the power of this assay. *T.b. brucei* infection resulted in significant increases in serum creatinine in both strains, compared to uninfected control mice (**Figure 1C**).

With evidence of azotaemia (elevated serum creatinine concentrations) and the propensity for *T.b. brucei* to invade tissues, we reasoned azotaemia might be a consequence of *T.b. brucei* invasion and colonisation of the kidney. We estimated the parasite burden in the kidneys using qPCR to quantify the *T.b. brucei*-specific *Pfr2* gene copies. We found that parasite DNA was detected in both mouse strains at both infection time points but not in the naïve controls. Susceptible BALB/c mice had higher parasite burden in the kidneys than tolerant C57BL/6 mice at both the early (1.6-fold) and late (3-fold) timepoints although this increase did not attain statistical significance (**Figure 1D**).

As our clinical and qPCR data indicated kidney injury in the presence of trypanosomes, we next evaluated the histopathology of the kidneys with a view to characterising the pattern of injury and detecting the location of the parasites.

To assess the impact of *T.b. brucei* infection on the structural integrity of the kidney, we examined H&E and PAS stained sections in each group of susceptible BALB/c and tolerant C57BL/6 (**Figure 2A and B**). Tubular degeneration and necrosis, which were the most prominent pathologic features, coincided with both mouse strain differences and time of infection. There were multiple foci of patchy tubular degeneration and necrosis at the early stage of infection in both mouse strains mostly in the outer stripe of the outer medulla. However, by the late time point (21 dpi), tubular necrosis was locally extensive and severe only in the susceptible BALB/c but not in the tolerant C57BL/6 mice. In the trypanotolerant C57BL/6 strain by day 21 of infection, there was evidence of regenerating tubules with dilated lumina and cytoplasmic basophilia of tubular epithelial cells were present. Inflammatory aggregates (comprised mostly of mononuclear cells) were present around the juxtamedullary region and peri-renal fat. Lesions in the glomeruli were evidenced by an increase in mesangial cells and an increased deposition of PAS-positive mesangial matrix in *T.b. brucei*-infected BALB/c) and C57BL/6 mice. These histologic findings are indicative of a mesangioproliferative glomerulopathy and consistent with a previous report (van Velthuysen and Florquin 2000), in addition to tubular injury and interstitial inflammation. Overall, the composite score of lesion severity was higher in BALB/c than C57BL/6 by day 21 of infection (*p* < 0.05) (**Table 1**).

**Figure 2.**
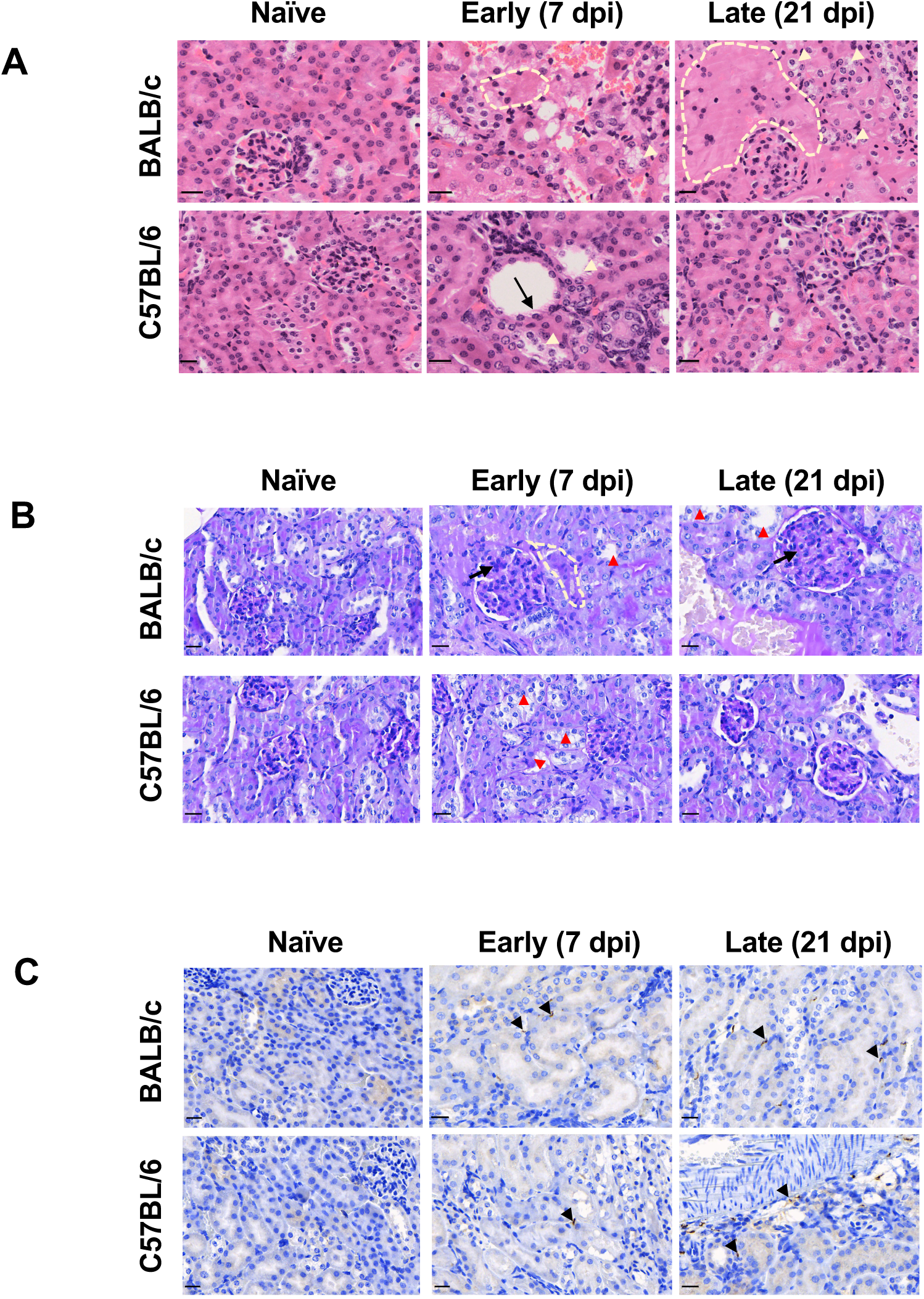
Trypanosoma brucei colonise the renal interstitium and elicits acute tubular injury in a temporal and strain-dependent manner. Kidney histopathology of trypanosusceptible BALB/c and trypanotolerant C57BL/6 mice with H&E (A) and PAS (B) staining. Bar: 20 µm. A) Intact tubules are present in naïve mice. There are multiple foci of vacuolar degeneration (yellow arrowheads) and simplification of epithelial cells of the proximal tubules (black arrow) during early stage infection (7 dpi). In addition, there are foci of acute tubular necrosis (broken outlines) in susceptible BALB/c which becomes progressively severe in the late time point (21 dpi) but attenuated in the tolerant C57BL/6. B) Tubular injury is evidenced by loss of PAS-positive brush border (red arrowheads). A focus of acute tubular necrosis (broken outline) is present in BALB/c mice at 7 dpi and there is proliferation of the mesangial cells within the glomeruli (black arrow) at both early and late timepoints that is absent in the tolerant C57BL/6 strain. C) Localisation of *Trypanosoma brucei* in the kidney of both susceptible (BALB/c) and tolerant (C57BL/6) mice (black arrowheads). Anti-trypanosome BIP antibody demonstrates that African trypanosomes are absent in naïve BALB/c and C57BL/6, but localise to the renal interstitium and outer medulla of both the susceptible (BALB/c) and tolerant (C57BL/6) mice as early as 7 dpi and remain at 21 dpi.

**Table 1.** Composite histopathologic score of mouse kidney. Ascending severity (scale of 0 - 9) of the kidney lesions during experimental *T.b. brucei* infection of relatively susceptible BALB/c and tolerant C57BL/6 mice.

| Infection Status | BALB/c<br>Median (n) | C57BL/6<br>Median (n) |
| --- | --- | --- |
| Naïve | 0 (3) | 0 (3) |
| Early [7 dpi] | 4 (3) | 5 (4) |
| Late [21 dpi]* | 6 (3) | 2 (3) |

\* Composite lesion severity score is significantly higher in BALB/c than C57BL/6 at this time point

We demonstrated the presence of *T.b. brucei* parasites in the kidney by immunohistochemistry staining with *T.b. brucei*-specific anti-BIP in infected mice. Extravascular parasites were observed in the kidney at both early (7 dpi) and late (21 dpi) in both strains of mice. Parasites were present and localised to the renal interstitium mostly in the outer medulla in infected mice of both strains (**Figure 2C)**. Our data demonstrate that *T.b. brucei* localises to the extravascular space in the murine kidney, in consonance with previous reports (De Niz *et al*. 2021; Mabille *et al*. 2022).

Thus, the pathologic findings indicate that trypanosomes occupy the kidney interstitium from at least the first peak of parasitaemia where it elicits patchy tubular damage accompanied by mononuclear inflammatory cellular aggregates in both strains, and suggestive of AKI-like phenotype. By late infection (21 dpi), pathologic findings diverged, with worsening tubular necrosis and a higher parasite burden in the susceptible BALB/c mice but a reparative tubular regeneration and lower parasite burden in the tolerant C57BL/6 strain. Taken together, the clinical and morphologic features observed in our models of *T. brucei* infection-induced kidney injury (TIKI) are consistent with AKI, and the pathology occurs in a mouse strain-dependent and temporal manner.

### Transcriptomic signature reflects clinicopathologic features of hypoxic AKI

The stark contrast in the progression of the pathology in both strains of mice from the early to late stages of infection raised the question of what molecular mechanisms underlie the temporal similarities and differences across both strains. To unravel this, we performed whole-kidney total mRNA sequencing of 3 – 4 biological replicates from BALB/c and C57BL/6 mice at both early (7 dpi) and late (21 dpi) time points and compared the differentially expressed genes (adjusted *p* < 0.05, log2fold change [LFC] either <0.5 or >0.5) relative to uninfected controls. Based on the top 500 most expressed genes, mouse strain (BALB/c or C57BL/6) and infection status (naïve, early and late infection) explained over 90% of the variance seen in the expression profiles (**Figure 3A).** In the sections that follow, we describe the transcriptomic findings in the context of known models of AKI using kidney-related differentially expressed genes, and the profound inflammatory and immune responses using functional gene analysis. Since we observed injury of the proximal tubules at histopathology, we compared our transcriptomic data to *a priori* list of genes specific to the proximal tubules in addition to known kidney injury markers (Liu *et al*. 2017).

**Figure 3.**
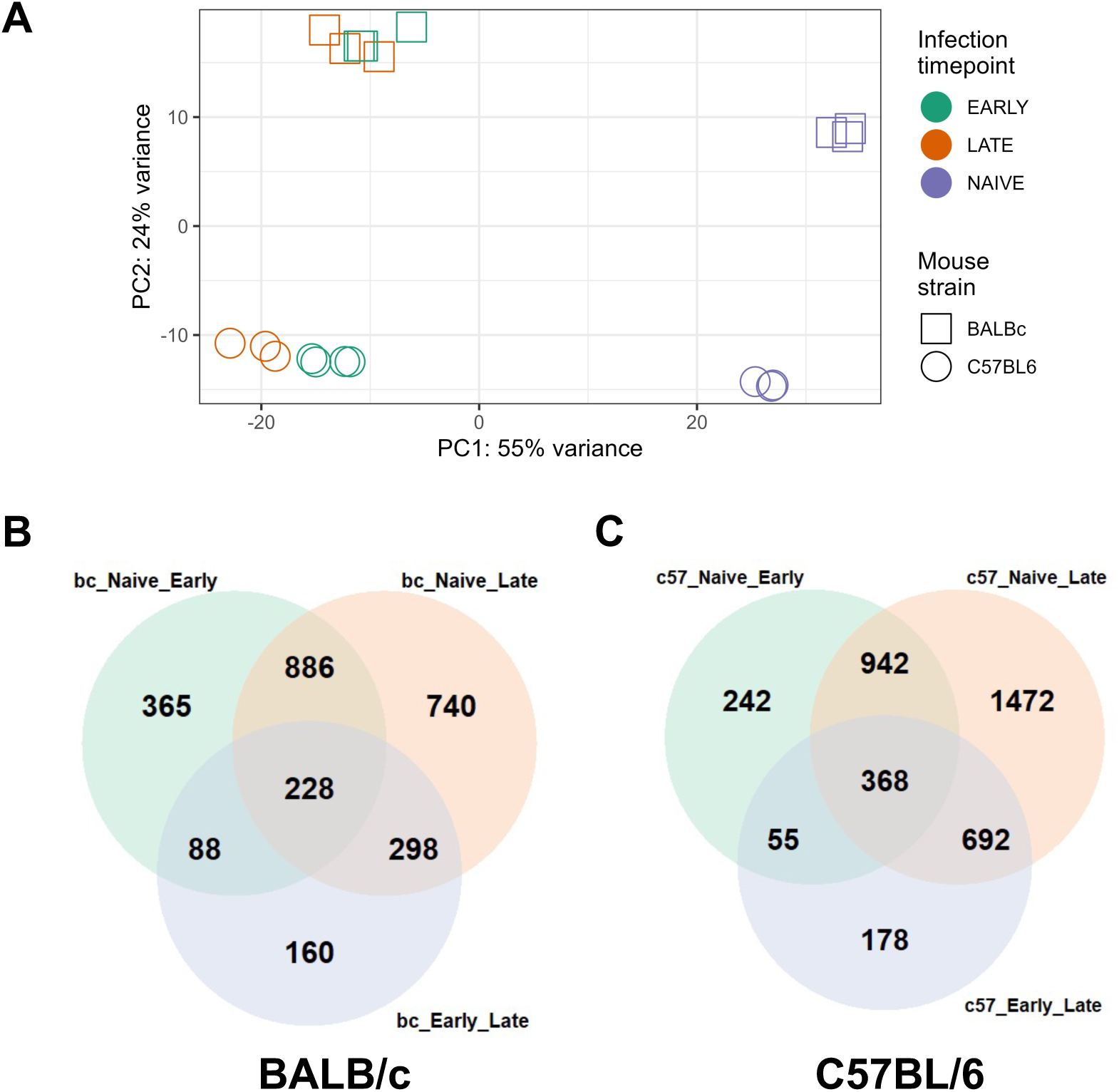
Differentially expressed genes from the kidneys during experimental *T. brucei* in susceptible BALB/c and tolerant C57BL/6 strains of mice. Principal component analysis plot (A) based on the top 1,000 most expressed genes. Mouse strain and infection status explain over 90% of variance between the groups. Differentially expressed genes in naïve versus early, naïve versus late, and early versus late comparison in BALB/c (B) and C57BL/6 (C) mice. Venn diagram shows the intersections and unique numbers of dysregulated genes (Log2FoldChange < -1 or >1, adjusted p < 0.05).

Consistent with the acute tubular degeneration and necrosis of proximal tubules, established kidney injury markers such as *Lcn2* and *Havcr1* (Liu *et al*. 2017; Gravina *et al*. 2023), were significantly elevated at both time points of infection in both mouse strains. Genes with endogenous expression in normal proximal tubules (*Ass1, Dio1, Miox*, *Slc7a13, Xpnpep2*) were downregulated (**Supplementary Table S2**). Of 67 solute carrier family genes (*Slc*) dysregulated in both strains at any time point during infection, 51 (76%) were downregulated. Notably, the sodium-independent organic anion transporters (solute carrier family 22) were downregulated. In addition, there was downregulation of pyruvate dehydrogenase (*Pdha1*), the enzyme that converts pyruvate to acetyl CoA (required for oxidative phosphorylation) in both mouse strains at both time points whereas both mouse strains during infection upregulated the glycolytic enzymes, hexokinase-3 (*Hk3*), phosphofructokinase platelet isoform (*Pfkp*) and pyruvate dehydrogenase kinase 3 (*Pdk3*) indicating a shift to glycolytic metabolism. We propose that the downregulation of genes involved in solute/ion transport and a metabolic shift to glycolysis is likely due to hypoxia resulting from the well reported infection-induced anaemia in mice (Amole *et al*. 1982; Stijlemans *et al*. 2018). Similarly, the patchy distribution of tubular necrosis is suggestive of a hypoxic/ischaemic insult. Although we did not measure erythrocyte indices clinically, we found that the canonical marker of hypoxia, carbonic anhydrase-9 (*Car9*) (Schaub *et al*. 2021), was markedly upregulated (relative to naïve controls) in both strains at early (BALB/c, LFC = 1.8; C57BL/6, LFC = 0.6) and late (BALB/c, LFC = 2.9; C57BL/6, LFC = 2.1) time points supporting an interpretation that TIKI is, at least in part, of hypoxic origin. Hypoxia was present early in infection and probably independent of the magnitude of parasitaemia (**Table 2**).

**Table 2.**
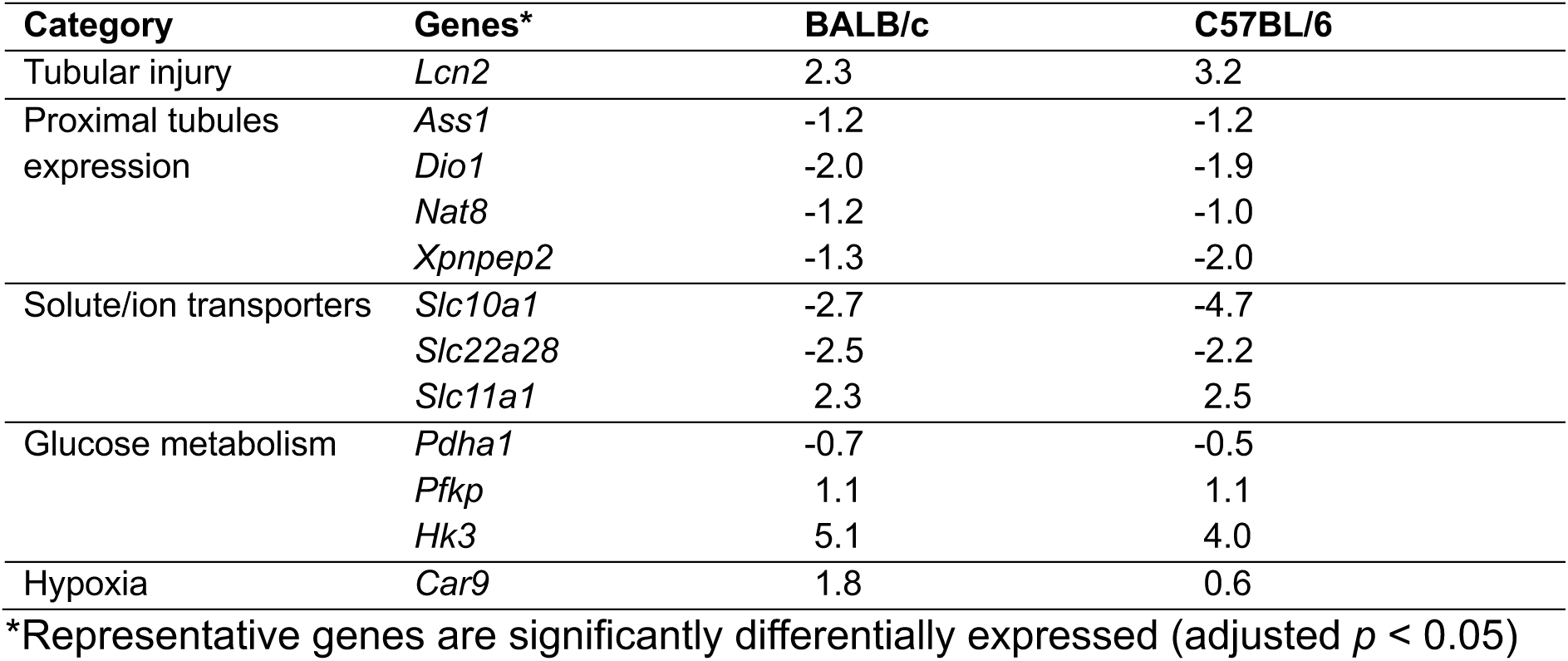
Tubule-specific expression of transcriptomic signature of *T. brucei*-induced hypoxic AKI at the early time of infection (7dpi)

To evaluate the response to hypoxia in both strains, we profiled the expression patterns of the hypoxia-inducible factor alpha (HIF-α) and downstream target genes. HIF-α orchestrates cellular adaptation to low oxygen tension, functioning as a master regulator of hundreds of genes in response to hypoxic conditions (Watts *et al*. 2020; Della Rocca *et al*. 2022). *Hif3α* was upregulated in both strains at 7 dpi (BALB/c, LFC = 1. 6; C57BL/6, LFC = 2.3), but only the trypanotolerant C57BL/6 strain by 21 dpi (LFC = 2.4). In the same vein, a downstream target gene of *Hif3α* involved in erythropoiesis, erythropoietin *(Epo),* was significantly upregulated only in C57BL/6 mice at both early (LFC = 4.5) and late (LFC = 5.9) time points but was not upregulated at any infection time point in BALB/c mice. Interestingly, other genes that function to increase oxygen delivery were upregulated in both mouse strains either at both infection time points (*Timp1*) or only at 21 dpi (*Nos2*, and *Hmox1*). Taken together, these findings suggest that recovery in renal structure from acute tubular injury seen in C57BL/6 might be linked to a more intense response to hypoxia including effective erythropoiesis and oxygen delivery to tissues.

### *T. brucei* elicits a phenotype consistent with other models of acute kidney injury (AKI) in mice

With evidence of morphologic and transcriptional features characteristic of AKI in both mouse strains at the early time point, we then compared the molecular features in our infection model with those reported in other models of AKI using a bioinformatic approach. We found that of the 212 genes that were common to four out of six models of AKI (Hultström *et al*. 2018), 120 genes (∼ 57%) were dysregulated (adjusted *p* < 0.05) at either early or late infection time point compared to naive. At the molecular level, AKI has been described as a stress response associated with active transcription, upregulation of genes involved in regeneration, apoptosis and survival, extracellular matrix organisation as well as a loss of mature phenotype (downregulation of genes that correspond to adult renal function) (Safirstein 2004). We found that while some transcription factors were upregulated (*Irf1*, *Irf7*, *Ddit3*, *Tgif1*, *Batf*) in both mouse strains, most were upregulated in C57BL/6 only and either downregulated (*Klf4, Klf6, Jund, Maff, Csrnp1*) or undetected (*Atf3*, *Elf4, Hif3a*) in BALB/c mice (**Table 3**). These strain-specific differential upregulation of factors might explain the reparative tubular response seen in C57BL/6 but not BALB/c mice at 21 dpi. Genes related to extracellular matrix organisation and involved in epithelial-mesenchymal transition (*Itgb2, Lgals3, Spock2, Tgfbi, Timp1, Mmp9, Mmp14*) were also upregulated (**Table 3**). These findings combined with the downregulation of genes involved in tubular function suggest that TIKI in mice show recapitulates the features seen in other models of AKI.

**Table 3.**
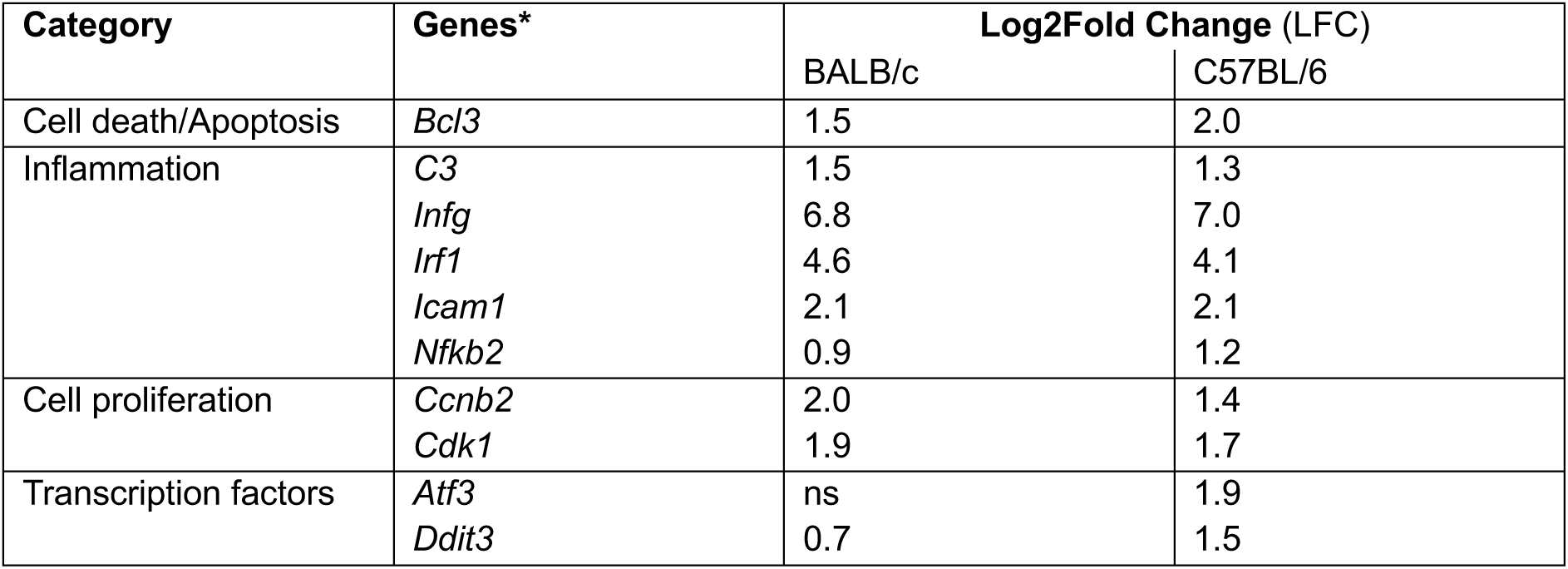
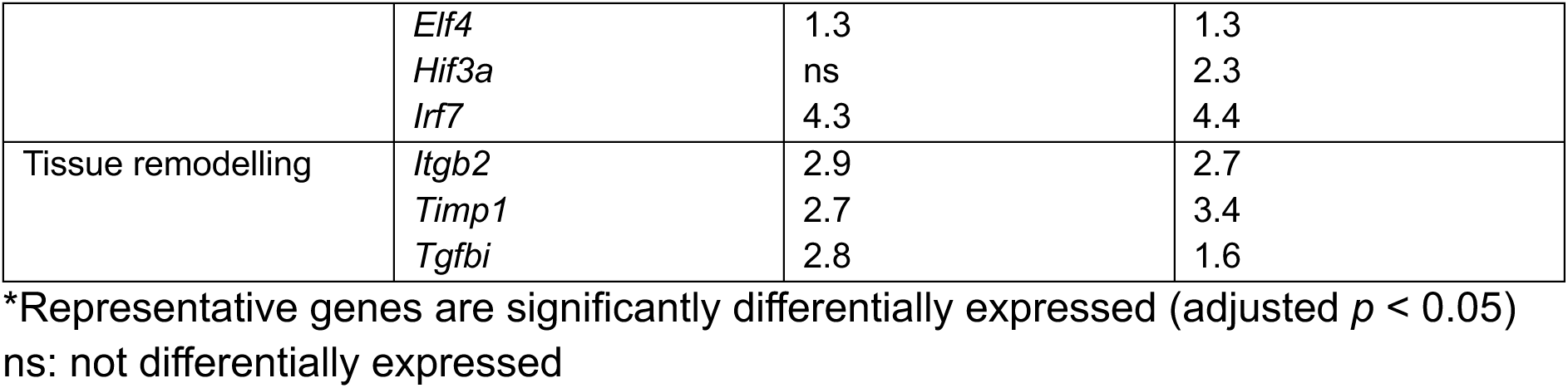
Common features of models of AKI with gene-specific expression at the early time of infection (7dpi) relative to uninfected controls differentially expressed

### *T. brucei* induces inflammatory and immune transcriptomic signatures in the kidneys

Following our histologic finding of inflammation during infection and in consonance with what has been previously reported, we characterised the transcriptomic landscape to gain broad insights into what mechanisms might be at play during infection by gene ontology analysis of lists of differentially expressed genes (DEGs; *adjusted p* < 0.05) whose log2fold-change (LFC) were either greater than 1 or less than -1. The number of DEGs for naïve-versus-early infection, naïve-versus-late infection, and early-versus-late infection comparisons were 1,567; 2,152 and 774, respectively, in susceptible BALB/c mice (**Figure 3B**). In the trypanotolerant C57BL/6 mice, the number of DEGs for naïve-versus-early infection, naïve-versus-late infection, and early-versus-late infection comparisons were 1,606; 3,473 and 1293 respectively (**Figure 3C**). Associated volcano plots for naïve-versus-early infection (**Figure 4A**), naïve-versus-late infection (**Figure 4B**) and early-versus-late infection (**Figure 4C**) in both mice strains are presented.

**Figure 4.**
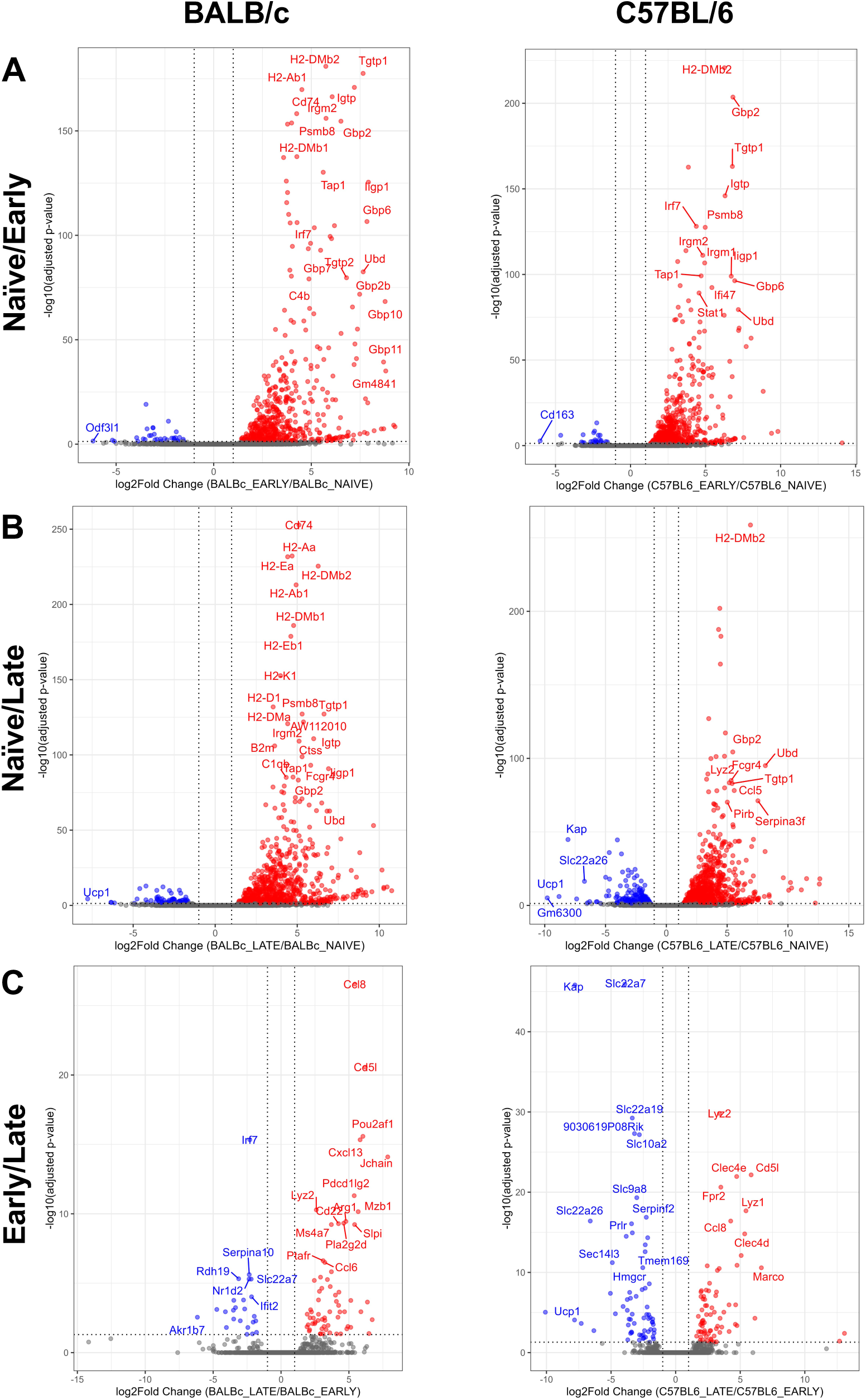
Volcano plots of differentially expressed genes in the kidneys during murine trypanosomiasis. Volcano plots of dysregulated genes indicated by upregulated (LFC >1 (red)) , downregulated (LFC < -1 (blue)) or not differentially expressed (LFC <1,>-1 (grey) genes for each timepoint comparison in each strain.

To get a clearer picture of the processes driven by the DEGs, we conducted gene ontology and functional analyses using gene lists derived from naïve-versus-early infection and naïve-versus-late infection comparisons, in both strains. The predominant downregulated biological processes included organic anion transport, monocarboxylic acid transport, cholesterol metabolism and fatty acid metabolism which were related to renal metabolism and function (**Table 4**). Functional gene analysis of upregulated genes (LFC > 1 & adjusted *p* < 0.05) in infected mice relative to naïve at both time points in both mouse strains revealed that the significantly enriched KEGG pathways were related to inflammation, phagocytosis, innate and adaptive immune responses, and haematopoiesis (**Figure 5A**). Thus, the clinicopathologic features of TIKI coincides with a transcriptional downregulation of renal tubular function, and in part mediated by the trypanosome-induced inflammatory/immune response as well as the hypoxia-driven tissue injury.

**Figure 5.**
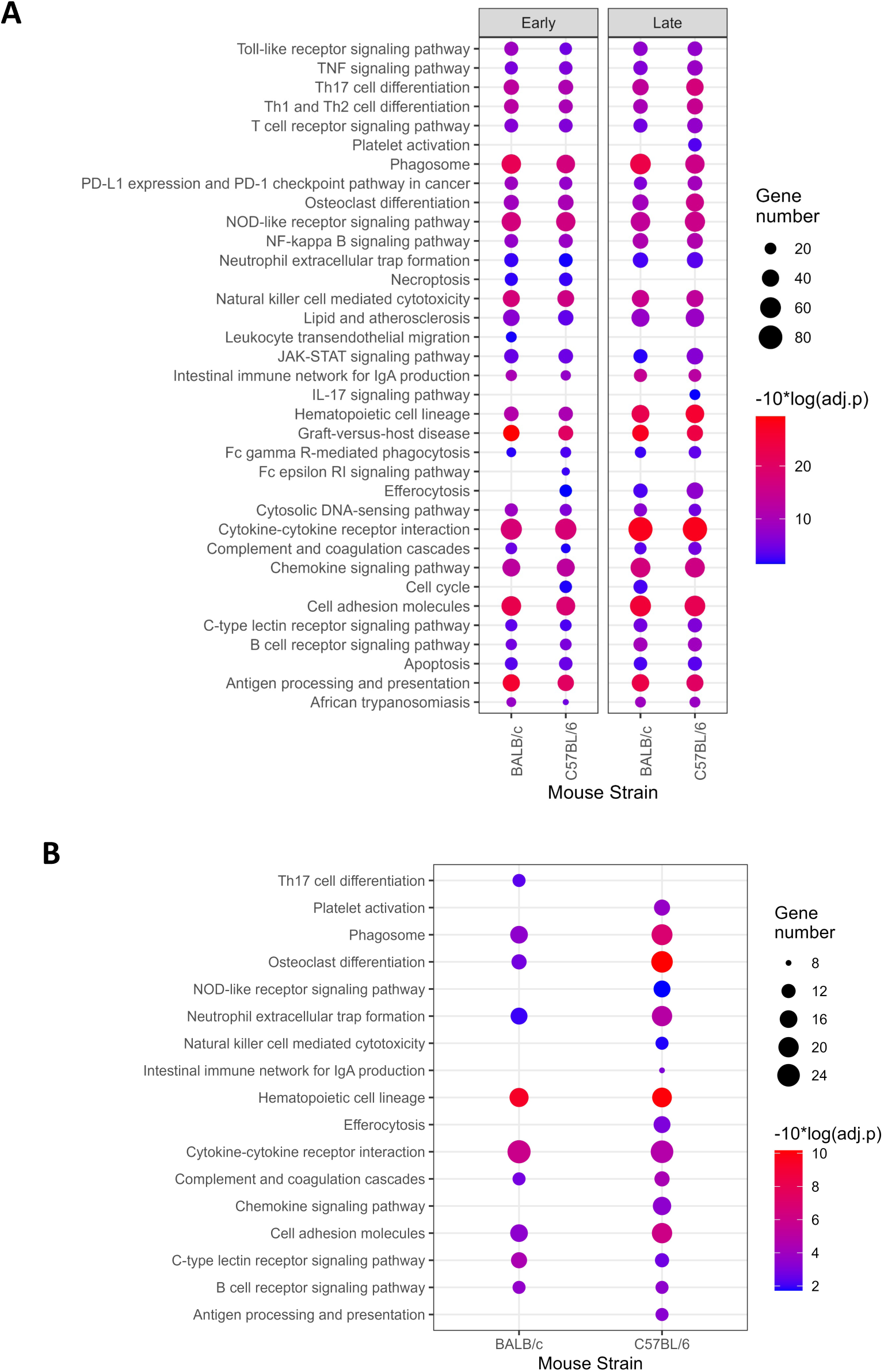
Transcriptome demonstrates broad immune and inflammatory response in the kidneys in a mouse strain-dependent manner. A) Enriched KEGG pathways across both susceptible BALB/c and tolerant C57BL/6 mice at either early (7 dpi) or late (21 dpi) timepoints in *Trypanosoma brucei* infection relative to naïve controls. B) Changes in Enriched KEGG pathways in the kidneys between early (7dpi) and late (21 dpi) time points in susceptible BALB/c and tolerant C57BL/6 mice infected with *T. brucei*. Size is number of genes and intensity is significance.

**Table 4.**
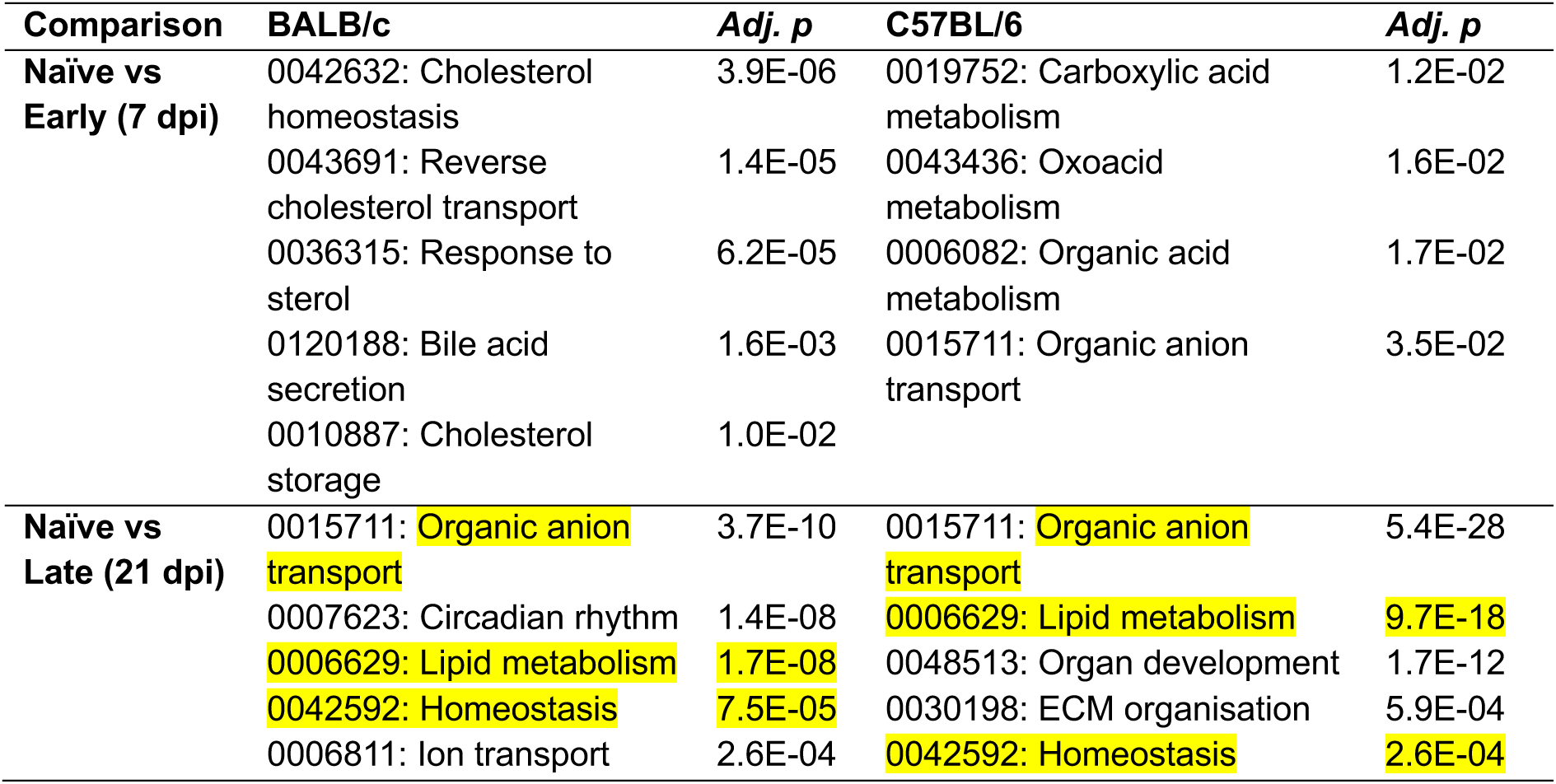
Top downregulated Gene Ontology-biological processes (GO-BP) in the kidney during experimental *T.b. brucei* infection of susceptible BALB/c and tolerant C57BL/6 mice relate mostly to ion transport. Common GO-BP terms between the two strains are highlighted in each comparison.

### Kidney damage is resolved in C57BL/6 but not BALB/c mice

Early in infection, we see a pattern of transcriptional changes that are consistent with the clinical and morphologic features of AKI, and these patterns are common to both strains of mice. However, we observed a profound difference in pathology in the mouse strains at the later time point. To investigate this further, we performed gene ontology on the genes which were differentially upregulated at the late stage, relative to early stage of infection in BALB/c and C57BL/6 mice. We identified enriched KEGG pathways that were common to both strains as well as those unique to each strain (**Figure 5B**).

Notably, only the Th17 signalling pathway was upregulated in susceptible BALB/c whereas KEGG pathways relating to innate immunity (platelet activation, chemokine signalling, antigen presentation, efferocytosis, natural killer cell mediated cytotoxicity and NOD-like receptor signalling) were upregulated in trypanotolerant C57BL/6 mice. Thus, whereas pathways regulating cell-mediated adaptive immunity were associated with susceptibility in BALB/c mice, pathways involving pathogen recognition, innate immunity and anti-inflammatory responses were associated with reduced organ pathology/and or improved tissue recovery in the tolerant C57BL/6. Consistent with the clinicopathology, the transcriptomic profiles of BALB/c and C57BL/6 mice by late infection (21 dpi), differ dramatically reflecting the worsening tubular necrosis in BALB/c mice but reparative tubular regeneration and lower parasite density in C57BL/6 mice.

### Cellular composition of the infected kidneys

To characterise the cellular compositions of the immune microenvironment of the kidneys, we performed *in silico* deconvolution of inflammatory cells population using the seq-ImmuCC pipeline (Chen *et al*. 2017). Our *in silico* deconvolution revealed stark strain differences. While cells of the mononuclear phagocyte system constituted at least 75% of immune cells and this proportion persisted throughout the course of infection in the susceptible BALB/c mice (**Figure 6A**), these cells accounted for only 50% in the tolerant C57BL/6. More intriguing is the finding of a massive expansion of CD8+ T cells early in the infection and the return to similar proportions as the naïve during late infection timepoint in C57BL/6 (**Figure 6B**). This suggests that CD8+ T cells might be important in parasite clearance in the tolerant C57BL/6 strain which is consistent with our observation of fewer parasites in the kidneys of these mice. Taken together, it would suggest that while a pro-inflammatory and cytotoxic response is required to clear parasites initially, milder tissue pathology is linked to effective clearance of parasites as well as timed control of the pro-inflammatory response.

**Figure 6.**
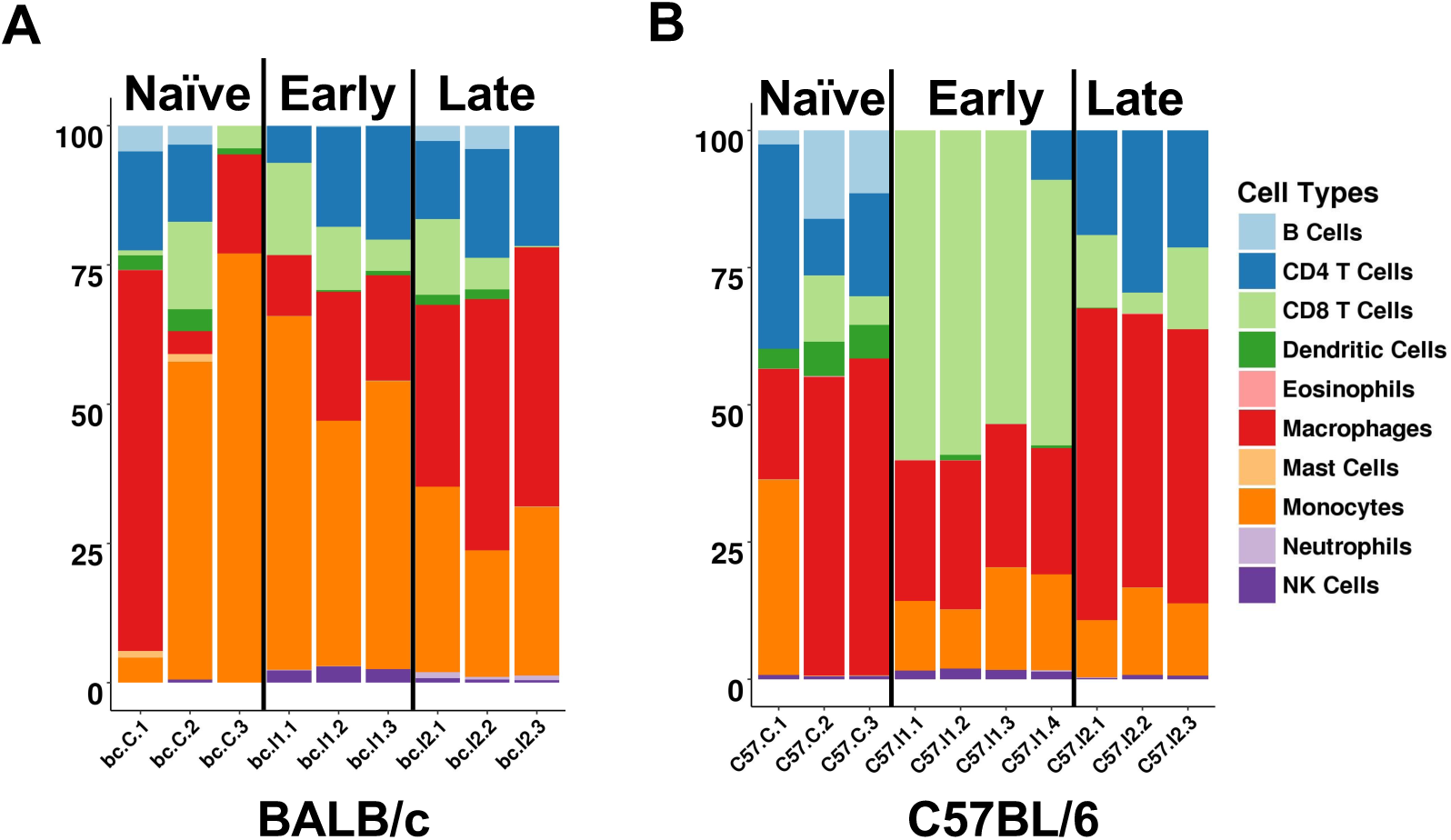
In silico deconvolution of immune cells proportions in the kidney. Proportions of immune cells are shown for each sample, for each timepoint, in each mouse strain for naïve and *T. brucei*-infected susceptible BALB/c (A) and tolerant C57BL/6 (B) mice.

### Serum creatinine is elevated in HAT cases

To confirm if the clinicopathologic features of murine TIKI was also observed in humans, we measured serum creatinine levels in trypanosome-infected individuals from the Democratic Republic of Congo. These serum samples, archived in the TrypanoGEN Biobank (Ilboudo *et al*. 2017), were from individuals who had active infections of *T.b. gambiense* as well as uninfected controls. We found that HAT cases had significantly elevated levels of serum creatinine compared to uninfected individuals (**Figure 7**). This finding indicates that similar mechanisms and renal responses might be at play in HAT as well as AAT, thus HAT cases carry the potential risk of developing TIKI, if not monitored.

**Figure 7.**
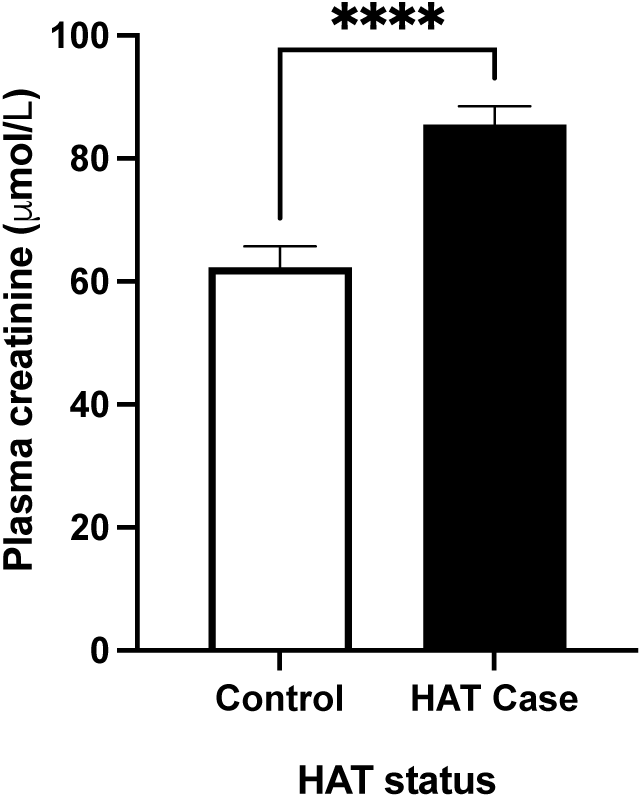
HAT cases from DRC show elevated plasma creatinine levels relative to endemic control individuals. Plasma creatinine levels were measured in humans infected with *T.b. gambiense* and uninfected individuals from HAT endemic foci in DRC. **** *p* < 0.0001.

## Discussion

In this work, we sought to characterise the histopathological and transcriptional responses of the kidney to *T. brucei* infection. We utilised well-defined murine models of trypanosusceptibility (BALB/c mice) and trypanotolerance (C57BL/6 mice) and evaluated the renal-specific host responses during the acute (7 dpi) and late (21 dpi) stages of infection. We demonstrate African trypanosomes in the renal interstitium, provide morphologic, and molecular evidence of acute tubular injury in a temporal- and strain-dependent manner, and identify strain-specific immune pathways upregulated throughout infection that likely mediate the differential renal pathology and host susceptibility. In addition, we found clinical evidence of significant elevation of serum creatinine in HAT patients and *T. brucei*-infected mice, thus showing TIKI as a model of AKI. Compared to other previously reported models of AKI, we demonstrated the similarities in the molecular responses that characterise TIKI.

The molecular basis of AKI appears to be the end-result of a conserved renal response to a variety of noxious stimuli. Previously described mouse models of AKI include ischaemia-reperfusion injury (transient clamping of renal artery), toxic injury (cisplatin), pigment nephropathy (following rhabdomyolysis or haemolysis), sepsis (caecal ligation and puncture) and obstructive injury (unilateral ureteral obstruction). Hultström *et al*. (2018) using an informatic approach, demonstrated that the transcriptomic signatures common to four different models of AKI included cell death, tissue remodelling, hypoxia, oxidative stress, and inflammation. In agreement with that report, we reported upregulation of genes related to cell injury and death (*Lcn2*, *Bcl3, Casp12*), hypoxia (*Car9*), tissue remodelling/extracellular matrix reorganisation (*Timp1*, *Plaur, Itgb2, Mmp14*) and interferon-driven inflammation (*Irf1, Ccl19, Cxcl10*) in both mouse strains. *T.b.brucei* infection induces anaemia (seen with ischaemic/hypoxic model), intravascular and extravascular haemolysis (pigment nephropathy model) as well as profound inflammation (sepsis-like model). This scenario more closely resembles what happens in humans in high-risk hospital settings: multifactorial concurrent renal insults.

Inflammation and the immune response contribute to both the elimination of the parasite as well as tissue pathology in host. Consistent with previous reports (Machado *et al*. 2021; Leigh *et al*. 2015), we report a lymphocytic inflammatory and immune response in the kidneys of both mice strains as early as 7 dpi. The strain-dependent differential morphologic features at the late time point of infection appears to be the result of a combination of the type of temporal immune responses, balance between pro-inflammatory and anti-inflammatory processes as well as an effective pro-survival transcriptional response to deleterious environmental cues. Our *in silico* deconvolution analysis revealed the marked expansion of CD8+ cytotoxic lymphocytes at early time point and a return to similar proportions as naïve by the late time point only in tolerant C57BL/6 mice. Compared to the susceptible BALB/c, the proportion of phagocytes were consistently elevated throughout infection. Maintaining a balance between pro-inflammatory and anti-inflammatory processes contributes to the differential severity of renal pathology. Notably, efferocytosis – a process which quells inflammation by stimulating the production of anti-inflammatory cytokines while simultaneously repressing proinflammatory cytokines (Doran *et al*. 2020), was upregulated between the early and late time point of infection only in the tolerant C57BL/6 mice. B cell expansion, which we report in both murine models, is a common feature with trypanosome infection (Magez *et al*. 2008; Quintana *et al*. 2022). We found that *Mcpip-1*, a critical negative regulator of inflammation which limits autoimmunity by its effect on B-cell expansion (Dobosz *et al*. 2021), is upregulated throughout infection only in tolerant C57BL/6 but not in susceptible BALB/c mice. Similarly, p21 (*Cdkn1a*) which has been reported to play a protective role in models of AKI (Safirstein 2004) was significantly upregulated only in C57BL/6 mice, coinciding with limited tissue pathology. Taken together, a fine control of the inflammatory and immune responses limits immunopathology.

Our models of relative resistance and susceptibility, recapitulate the well characterised clinical features associated with *T. brucei* infection (Magez and Caljon 2011; Naessens 2006). Both mouse strains develop similar initial peaks of parasitaemia early in infection. Whereas tolerant mice clear the initial and subsequent peaks of parasitaemia to almost undetectable levels, susceptible mice do not. The severity of renal pathology and associated inflammation mirrored this trend: initial tubular injury in both mouse strains at the early time point, but recovery in tolerant mice and worsening injury in susceptible mice at the late time point. However, clinical (serum creatinine) and molecular (*Lcn2*) markers of AKI did not mirror the recovery in structure seen by histology in the tolerant C57BL/6. We cannot rule out that the elevated serum creatinine is not due in part to other pre-renal factors such as dehydration, although clinical signs, for example loss of skin elasticity, were not observed during the infection. It is likely that restoration of structure precedes the restoration of normal function in the kidney.

Over the last decade, a link between immunity against HAT and the risk of developing chronic kidney disease has been identified for carriers of recessive risk alleles in the primate-specific apolipoprotein L1 (APOL1) gene (Genovese et al. 2010; Cooper et al. 2017). Studies that demonstrate this heightened risk of CKD have been elucidated mostly in people outside of the tsetse-endemic, sub-Saharan parts of the continent (Genovese *et al*. 2010; Tzur *et al*. 2010). Currently, HAT patients are not routinely screened for markers of renal function. Our results show that HAT cases may be at risk of azotaemia and thus AKI. Untreated AKI can itself predispose to CKD (Liu *et al*. 2017). Our murine model of TIKI presented here demonstrates the effects of trypanosomes on kidneys in the absence of APOL1 and represent a tool to understand the effect of African trypanosomes on the host’s renal function especially in long-term infections. The contribution of HAT to the development of AKI and subsequently CKD in at-risk populations remains undetermined.

## Supplementary Tables Legend

**Supplementary Table S1**: Read counts and percentage uniquely mapped from kidney RNA sequencing analysis.

**Supplementary Table S2:** Read counts for genes with endogenous expression in the proximal convoluted tubules (PCT).

## Notes

### Competing Interest Statement

The authors have declared no competing interest.

