## supplemental file 1 for "Murine trypanosomiasis recapitulates transcriptomic features of acute kidney injury"

**Supplemental Table 1: Percentage Mapping of Reads**

| Sample name | Mouse strain | Infection Status | Input Reads | Uniquely Mapped | Mapping % |
|---|---|---|---|---|---|
| B6_i11 | C57BL/6 | 7 dpi | 32700895 | 27181418 | 83% |
| B6_i12 | C57BL/6 | 7 dpi | 32514849 | 26894363 | 83% |
| B6_i13 | C57BL/6 | 7 dpi | 33603301 | 27822841 | 83% |
| B6_i14 | C57BL/6 | 7 dpi | 33624514 | 27908894 | 83% |
| B6_i21 | C57BL/6 | 21 dpi | 32579218 | 26902793 | 83% |
| B6_i22 | C57BL/6 | 21 dpi | 32720223 | 27503162 | 84% |
| B6_i23 | C57BL/6 | 21 dpi | 33343375 | 27696135 | 83% |
| B6_N1 | C57BL/6 | NAÏVE | 32817007 | 27448336 | 84% |
| B6_N2 | C57BL/6 | NAÏVE | 34067311 | 28813331 | 85% |
| B6_N3 | C57BL/6 | NAÏVE | 33660715 | 27692258 | 82% |
| Bc_21 | BALB/c | 21 dpi | 33325831 | 27932344 | 84% |
| Bc_22 | BALB/c | 21 dpi | 33043129 | 28351456 | 86% |
| Bc_23 | BALB/c | 21 dpi | 33337403 | 27714940 | 83% |
| Bc_i11 | BALB/c | 7 dpi | 33035476 | 27217776 | 82% |
| Bc_i12 | BALB/c | 7 dpi | 32715552 | 26681412 | 82% |
| Bc_i13 | BALB/c | 7 dpi | 33028688 | 27477213 | 83% |
| Bc_N1 | BALB/c | NAÏVE | 33544453 | 28492589 | 85% |
| Bc_N2 | BALB/c | NAÏVE | 33566113 | 27988750 | 83% |
| Bc_N3 | BALB/c | NAÏVE | 32787159 | 27987004 | 85% |
