## supplemental file 2 for "Murine trypanosomiasis recapitulates transcriptomic features of acute kidney injury"

**Supplemental Table 2: Read Counts for genes with PCT-specific expression**

| Sample name | B6_i11 | B6_i12 | B6_i13 | B6_i14 | B6_i21 | B6_i22 | B6_i23 | B6_N1 | B6_N2 | B6_N3 | Bc_21 | Bc_22 |
|---|---|---|---|---|---|---|---|---|---|---|---|---|
| Mouse strain | C57BL/6 | C57BL/6 | C57BL/6 | C57BL/6 | C57BL/6 | C57BL/6 | C57BL/6 | C57BL/6 | C57BL/6 | C57BL/6 | BALB/c | BALB/c |
| Infection Status | 7 dpi | 7 dpi | 7 dpi | 7 dpi | 21 dpi | 21 dpi | 21 dpi | NAÏVE | NAÏVE | NAÏVE | 21 dpi | 21 dpi |
| Ass1 | 28921 | 28885 | 33770 | 44154 | 54328 | 77460 | 46348 | 87664 | 69072 | 66246 | 46505 | 49760 |
| Fbp1 | 14960 | 13136 | 15958 | 18483 | 11910 | 19049 | 15699 | 24024 | 25058 | 22576 | 26972 | 24657 |
| Miox | 26349 | 32271 | 31229 | 35077 | 26258 | 34330 | 24724 | 80178 | 89306 | 71965 | 36335 | 38440 |
| Napsa | 50015 | 44982 | 52943 | 56354 | 30921 | 50040 | 49009 | 40954 | 56594 | 42469 | 40753 | 40142 |
| Cyp2e1 | 1837 | 2610 | 1593 | 2893 | 2564 | 1353 | 1736 | 4867 | 3469 | 6397 | 1922 | 6729 |
| Slc22a18 | 6923 | 7451 | 6978 | 10059 | 6791 | 8274 | 7138 | 10828 | 12951 | 10948 | 6328 | 5822 |
| Anxa13 | 28 | 27 | 45 | 30 | 27 | 41 | 25 | 55 | 41 | 86 | 19 | 12 |
| Acss2 | 3111 | 3484 | 3389 | 3940 | 2366 | 2661 | 2634 | 5400 | 4606 | 4558 | 3003 | 3035 |
| Aspg | 439 | 394 | 390 | 541 | 178 | 232 | 254 | 600 | 586 | 579 | 567 | 322 |
| Thnsl2 | 4213 | 4197 | 4869 | 5344 | 3636 | 4791 | 4473 | 5054 | 5173 | 4992 | 4848 | 4008 |
| Dmgdh | 1786 | 1480 | 1826 | 2112 | 750 | 1190 | 1127 | 2010 | 2916 | 2698 | 2183 | 1699 |
| Afm | 816 | 718 | 936 | 741 | 550 | 791 | 729 | 1020 | 1084 | 1205 | 1136 | 1115 |
| Inmt | 679 | 1075 | 724 | 1548 | 880 | 2021 | 1012 | 4626 | 4444 | 4712 | 269 | 193 |
| Aspdh | 931 | 1007 | 981 | 1427 | 444 | 714 | 652 | 1303 | 1636 | 1374 | 1201 | 859 |
| Dpys | 230 | 292 | 262 | 324 | 66 | 116 | 94 | 474 | 571 | 488 | 237 | 194 |
| Pyroxd2 | 1833 | 2074 | 2333 | 2427 | 2243 | 2726 | 2223 | 3663 | 3513 | 3557 | 2062 | 1979 |
| Eci3 | 2414 | 2365 | 2616 | 2461 | 1057 | 1935 | 1799 | 3306 | 3819 | 3709 | 1506 | 1491 |
| Acy3 | 27500 | 25448 | 24998 | 35200 | 12932 | 20713 | 20363 | 25876 | 30086 | 24614 | 24710 | 21727 |
| Nat8 | 751 | 462 | 544 | 797 | 1047 | 1101 | 919 | 1271 | 1369 | 1104 | 1242 | 1096 |
| Kap | 180334 | 222898 | 225871 | 208859 | 392 | 459 | 2034 | 221757 | 267911 | 257788 | 14566 | 13163 |
| Slc7a13 | 21 | 23 | 21 | 45 | 0 | 8 | 5 | 81 | 40 | 47 | 0 | 5 |
| Pm20d1 | 850 | 813 | 885 | 949 | 674 | 764 | 653 | 1519 | 1533 | 1605 | 4458 | 3781 |
| Apom | 1053 | 917 | 935 | 1427 | 1134 | 1286 | 1107 | 2246 | 1776 | 1608 | 1407 | 1175 |
| Haao | 2367 | 2105 | 2456 | 2801 | 2226 | 2826 | 2695 | 2988 | 3857 | 3390 | 3135 | 2702 |
| Khk | 10237 | 9224 | 11035 | 12699 | 8844 | 10919 | 10446 | 11294 | 14759 | 13254 | 13702 | 12233 |
| Upb1 | 2601 | 2350 | 2915 | 2964 | 2597 | 3983 | 3117 | 3215 | 3274 | 3399 | 3009 | 3153 |
| Amacr | 1913 | 2047 | 1996 | 2559 | 1423 | 1819 | 1778 | 2389 | 2542 | 2241 | 2384 | 2520 |
| Bhmt2 | 1801 | 1539 | 1651 | 1988 | 889 | 1202 | 1311 | 1740 | 2263 | 2210 | 2301 | 1993 |
| Hsd11b1 | 1545 | 2186 | 1532 | 1343 | 2032 | 1432 | 1709 | 1142 | 1465 | 1545 | 1620 | 980 |
| Lrp2 | 35548 | 34305 | 43169 | 31639 | 14500 | 22789 | 20539 | 56199 | 52077 | 61098 | 30260 | 30383 |
| Mep1a | 24069 | 25072 | 32265 | 29085 | 9756 | 13401 | 13216 | 32977 | 42005 | 36191 | 14667 | 14590 |
| Dio1 | 481 | 560 | 650 | 674 | 119 | 218 | 255 | 1552 | 2572 | 2605 | 156 | 85 |
| Proz | 49 | 73 | 79 | 65 | 63 | 46 | 54 | 173 | 185 | 170 | 65 | 48 |
| Xpnpep2 | 173 | 147 | 162 | 221 | 437 | 526 | 589 | 629 | 797 | 659 | 621 | 695 |

**Supplemental Table 2: Read Counts for genes with PCT-specific expression**

| Sample name | Bc_23 | Bc_i11 | Bc_i12 | Bc_i13 | Bc_N1 | Bc_N2 | Bc_N3 |
|---|---|---|---|---|---|---|---|
| Mouse strain | BALB/c | BALB/c | BALB/c | BALB/c | BALB/c | BALB/c | BALB/c |
| DPI group | 21 dpi | 7 dpi | 7 dpi | 7 dpi | NAÏVE | NAÏVE | NAÏVE |
| Ass1 | 52087 | 26458 | 32512 | 31832 | 58099 | 62182 | 62199 |
| Fbp1 | 21633 | 20165 | 19901 | 21537 | 24899 | 24218 | 29422 |
| Miox | 42881 | 20388 | 35083 | 33047 | 58555 | 70442 | 61538 |
| Napsa | 38859 | 63142 | 59249 | 53290 | 40029 | 46958 | 48642 |
| Cyp2e1 | 1404 | 7703 | 2769 | 2550 | 5069 | 3701 | 8829 |
| Slc22a18 | 6959 | 5490 | 6574 | 5571 | 7798 | 8888 | 6793 |
| Anxa13 | 15 | 3 | 2 | 8 | 11 | 7 | 12 |
| Acss2 | 3186 | 3759 | 4150 | 4044 | 4531 | 4146 | 4347 |
| Aspg | 488 | 904 | 889 | 761 | 605 | 412 | 481 |
| Thnsl2 | 4430 | 5311 | 5789 | 5811 | 4593 | 4142 | 5370 |
| Dmgdh | 1964 | 2441 | 2171 | 2283 | 2723 | 2647 | 2502 |
| Afm | 1171 | 878 | 1296 | 1077 | 1323 | 1496 | 1210 |
| Inmt | 315 | 501 | 251 | 415 | 725 | 834 | 840 |
| Aspdh | 1787 | 1378 | 1374 | 1542 | 1909 | 1894 | 1859 |
| Dpys | 258 | 109 | 110 | 155 | 364 | 550 | 279 |
| Pyroxd2 | 2105 | 1859 | 1385 | 1481 | 1702 | 2237 | 2114 |
| Eci3 | 1562 | 1249 | 1632 | 1741 | 1913 | 1970 | 2128 |
| Acy3 | 31130 | 37425 | 54395 | 50907 | 32764 | 30841 | 31357 |
| Nat8 | 1082 | 1357 | 1003 | 1004 | 2178 | 2077 | 2493 |
| Kap | 7789 | 154703 | 124632 | 130360 | 132646 | 108575 | 144726 |
| Slc7a13 | 0 | 3 | 14 | 8 | 16 | 30 | 34 |
| Pm20d1 | 3864 | 4679 | 4634 | 4403 | 6395 | 6874 | 5638 |
| Apom | 1144 | 1102 | 1305 | 1145 | 1689 | 1774 | 1698 |
| Haao | 3502 | 2778 | 2671 | 2646 | 3331 | 3585 | 3443 |
| Khk | 12490 | 12529 | 11811 | 11230 | 12495 | 14879 | 13544 |
| Upb1 | 2949 | 3502 | 2836 | 2998 | 3271 | 3135 | 2781 |
| Amacr | 2592 | 2260 | 2755 | 2549 | 2331 | 2364 | 2699 |
| Bhmt2 | 2403 | 2682 | 2214 | 2007 | 2452 | 2132 | 2147 |
| Hsd11b1 | 1109 | 1145 | 916 | 1336 | 936 | 923 | 1560 |
| Lrp2 | 29806 | 34341 | 41623 | 51705 | 47535 | 40951 | 49493 |
| Mep1a | 12740 | 21912 | 22761 | 24478 | 24385 | 24207 | 22545 |
| Dio1 | 107 | 173 | 229 | 204 | 724 | 755 | 744 |
| Proz | 65 | 65 | 69 | 46 | 99 | 117 | 110 |
| Xpnpep2 | 631 | 323 | 462 | 326 | 752 | 872 | 739 |
